# TreeToReads - a pipeline for simulating raw reads from phylogenies

**DOI:** 10.1101/037655

**Authors:** Emily Jane McTavish, James Pettengill, Steve Davis, Hugh Rand, Errol Strain, Marc Allard, Ruth E. Timme

**Affiliations:** Department of Ecology and Evolutionary Biology, University of Kansas, Lawrence KS, USA; Center for Food Safety and Nutrition, Food and Drug Administration, College Park, MD

## Abstract

Using genome-wide SNP-based methods for tracking pathogens has become standard practice in academia and public health agencies. There are multiple computational approaches available that perform a similar task: call variants by mapping short read data against a reference genome, quality filter these variants, then concatenate the variants into a sequence matrix for downstream phylogenetic analysis. However, there are no existing methods to validate the accuracy of these approaches despite the fact that we know there are parameters that can affect whether a SNP is called, or the correct tree is recovered. We present a simulation approach (TreeToReads) to generate raw read data from mutated genomes simulated under a known phylogeny. The user can vary parameters of interest at each step in the simulation (e.g., topology, model of sequence evolution, and read coverage) to assess the robustness of a given result, which is critical within both research and applied settings. Source code, examples, and a tutorial are available at https://github.com/snacktavish/TreeToReads.

## 1 Introduction

Genomics has revolutionized our understanding of patterns and processes of evolution across a wide range of taxa. Differentiating among individuals who only very recently diverged, between which only a few single nucleotide polymorphisms (SNPs) may exist, is only possible in the light of whole genome sequence data. In lieu of whole genome alignment, analysis pipelines attempt to extract the variable sites directly from the raw sequence reads (e.g. Illumina MiSeq data) and then infer the phylogeny directly from a SNP matrix of variable-only sites. In these examples where estimates of ancestry rely on a handful of data points, it is particularly important to ensure that analysis methods are validated and free from bias. Rigorous testing of these methods is needed, especially when the phylogenetic trees are used by public heath agencies to make regulatory decisions (e.g. identifying a source in a foodborne outbreak (*Hoffmann et al*., 2015).

**Figure 1:**
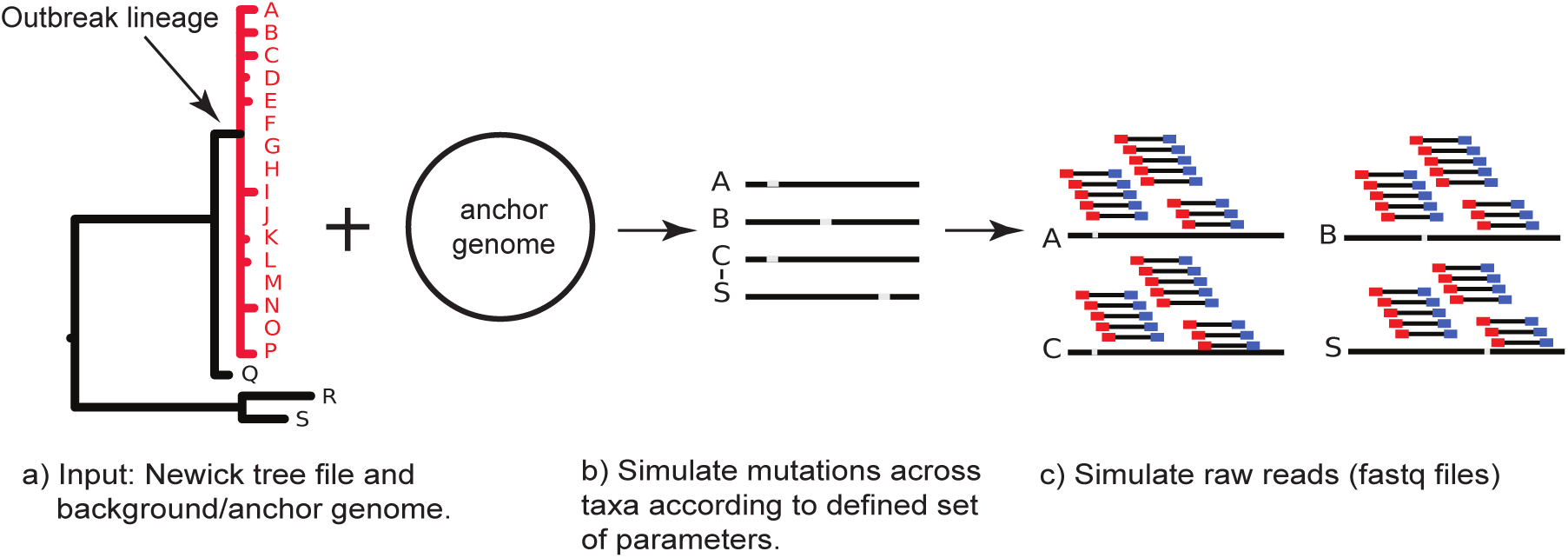
Schematic of the TreeToReads procedure

SNP-calling biases can be caused by various factors, including genotyping of single nucleotides which are polymorphic in a subset of the population (McTavish and Hillis, 2015), missing data cutoffs resulting in preferential inclusion of loci evolving at lower rates (Huang and Knowles, 2014) and those related to read mapping due to choice of reference genome (Bertels *et al*., 2014)). filter artifacts (Li, 2014) and different mapping algorithms (Pightling *et al*., 2014). Mis-estimation can be exacerbated by interaction between these dataset biases and analysis choices; for example using a model of evolution developed for sequence data on a panel of exclusively variable sites (Lewis, 2001) or choosing an inappropriate model of evolution (Sullivan and Swofford, 1997). Despite the sheer quantities of genomic data, it is possible that these types of biases could affect phylogenetic conclusions and, if systematic, inappropriate methods may converge to an incorrect result with high bootstrap confidence. In order to adopt data analysis pipelines for the regulatory environment it is necessary to understand these biases and validate their use. Without in silico modeling food safety scientists would have to rely on benchmark datasets where the truth can never be truly known.

Here, we present TreeToReads (TTR), a software tool to simulate realistic patterns of sequence variation across phylogenies in order to assess the robustness of evolutionary inferences from whole genome data to potential biases in the data collection and analysis pipeline.

## 2 Methods

The TTR pipeline generates short read data from genomes simulated along an input phylogeny. The software is written in python and requires two input files - a phylogeny with branch lengths and an anchor genome (Figure 1a); there is also a default configuration file within which the user can specify parameter settings (e.g., number of variable sites to simulate and nucleotide substitution model parameters). The branch lengths of the user provided phylogeny determine the probability that a single site is affected by multiple mutational events (Sukumaran and Holder, 2010). The pipeline uses seq-gen (Rambaut and Grass, 1997) to simulate the variable sites specified in the configuration file. These sites are then distributed across the anchor genome (Figure 1b). The locations for mutations either drawn from a uniform distribution, or clustered according to parameters of an exponential distribution specified in the configuration file. This procedure creates an output folder for each tip in the tree that contains the simulated genomes (fasta files). Using these simulated genomes, TTR calls the read simulation software, ART (Huang *et al*., 2012) to generate Illumina MiSeq paired-end reads (Figure 1c). The user can specify a different sequence error model in the configuration file. While TTR currently only supports automated generation of Illumina paired end reads, the simulated genome files may be used outside of TTR with any ART parameter configuration. Alternatively, if RAD-seq like data are desired other raw-read generators such as SimRAD (Lepais and Weir, 2014) can be used. If ART is invoked in TTR the program will output a fastq folder containing directories labeled with the names of each tip from the simulation tree within which the simulated reads in .fastq.gz and .sam format are deposited. A file with the location and nucleotide state of each mutation within each tip is also provided.

## 3 Case Study

To illustrate the utility of TTR, we tested the effects of sequence coverage on the ability of the CFSAN SNP pipeline (Davis *et al*., 2015) to call SNPs and recover an observed phylogeny (more importantly, the outbreak clade) (Figure 1a) of ten Salmonella enterica subsp. enterica serovar Bareilly sequences associated with a 2010 outbreak (Hoffmann *et al*., 2015). For outbreak regulatory decisions the most important part of a foodborne outbreak phylogenetic tree is the split that separates isolates belonging to the outbreak versus not part of the outbreak. We used a closed Salmonella enterica genome (CFSAN000189, GenBank: CP006053.1) as the anchor, simulated 150 variable sites under the GTR model, SNP clustering ON with locations for 20% drawing from an exponential distribution with a 125bp mean, and finally, a read error profile based on observed data. TTR was run under four different sequence coverage settings: 1X, 5X, 15X, and 30X. We analyzed the resulting four short read datasets with the CFSAN SNP pipeline and default settings (Davis *et al*., 2015), which identified SNPS within each set. Finally, we inferred the phylogeny for each set using RAxML (Stamatakis, 2014) under the ASCGTRCAT model. Results are as follows: 1X) zero SNPS, no phylogeny; 5X) 37 SNPS, correct outbreak (OB) clade; 15X) 146 SNPS, correct OB clade; 30X) 148 SNPS, correct OB clade. While this is a very simple test case, it illustrates the utility of TTR to test important parameters and their interactions affecting analysis pipelines ability to accurately call SNPS and infer phylogenies.

## 4 Conclusions

To date the phylogenetic perspective in simulation testing of assembly and alignment tools has been lacking in genomic simulation software. TreeToReads allows researchers to test the joint effects of multiple parameter values (such as coverage thresholds, amount of sequence variation, choice of reference genome, phylogenetic inference method, etc.) on the ability of any analysis pipeline to recover the signal and infer the correct tree. Simulating data that spans these parameters will help validate methods for reconstructing phylogenies directly from short-read data, which is especially useful for public health agencies using these methods to track various emerging pathogens.

